# Quantitative analysis of ZFY and CTCF reveals dependent recognition of tandem zinc finger proteins

**DOI:** 10.1101/637298

**Authors:** Zheng Zuo, Timothy Billings, Michael Walker, Petko M. Petkov, Gary D. Stormo, Polly M. Fordyce

## Abstract

The human genome contains around 800 C2H2 Zinc Finger Proteins (ZFPs), and many of them are composed of long tandem arrays of zinc fingers. Current motif prediction models assume longer finger arrays correspond to longer DNA-binding motifs and higher specificity. However, recent experimental efforts to identify ZFP binding sites *in vivo* contradict this assumption, with many having short motifs. Here, we systematically test how multiple zinc fingers contribute to binding for three model ZFPs: Zinc Finger Y (ZFY), CTCF, and ZNF343. Using ZFY, which contains 13 fingers, we quantitatively characterize its binding specificity with several methods, including Affinity-seq, HT-SELEX, Spec-seq and fluorescence anisotropy, and find evidence for ‘dependent recognition’ where downstream fingers can recognize some extended motifs only in the presence of an intact core site. For the genomic insulator CTCF, additional high-throughput affinity measurements reveal that its upstream specificity profile depends on the strength of the core, violating presumed additivity and position-independence. Moreover, the effect of different epigenetic modifications within the core site depends on the strength of flanking upstream site, providing new insight into how the previously identified intellectual disability-causing and cancer-related mutant R567W disrupts upstream recognition and deregulates CTCF’s methylation sensitivity. Lastly, we used ZNF343 as example to show that a simple iterative motif analysis strategy based on a small set of prefixed cores can reveal the dependent relationship between cores and upstream motifs. These results establish that the current underestimation of ZFPs motif lengths is due to our lack of understanding of intrinsic properties of tandem zinc finger recognition, including irregular motif structure, variable spacing, and dependent recognition between sub-motifs. These results also motivate a need for better recognition models beyond additive, position-weight matrix to predict ZFP specificities, occupancies, and the molecular mechanisms of disease mutations.

## Introduction

The zinc finger domain was first described in the TFIIIA protein of Xenopus which contains an array of nine zinc fingers involved in the recognition of RNA Polymerase III promoters^1,2^. Since then, it has been discovered that ZFPs occur in all eukaryotic species and have expanded enormously in the vertebrate lineage^3–5^ where they are the most abundant class of transcription factors (TFs)^6,7^. While ZFPs can have other roles, such as binding to RNA and in protein-protein interactions, it is generally thought that C2H2 family ZFPs function as DNA-binding TFs. Most C2H2 family ZFPs are composed of tandem arrays of zinc fingers, and the number of fingers for each ZFP has greatly expanded in vertebrates, with some human ZFPs containing more than 30 fingers (Fig. 1A).

**Figure 1.**
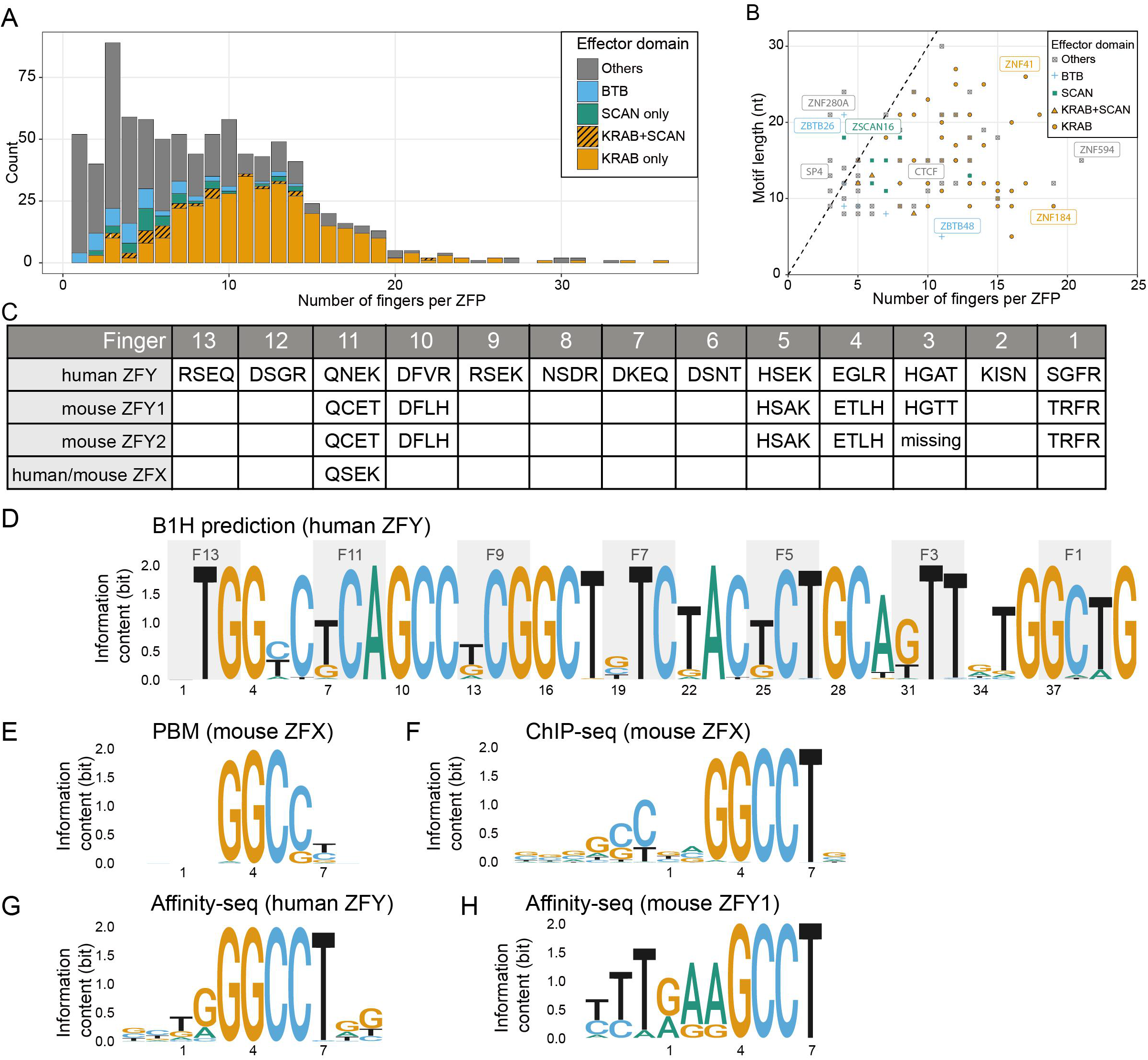
The statistics of human ZFPs and currently identified short motifs of ZFY and its orthologs. **A)** Distribution of C2H2-type zinc finger proteins (ZFPs) in human genome; **B)** Currently identified motif length vs. the number of fingers within each ZFP; **C)** Contact residues of Zinc Finger Y (ZFY) and its close orthlogs, mZFY1, mZFY2, and ZFX; **D)** Predicted motif by B1H method; **E)** Motif of mouse ZFX identified by protein binding microarray (PBM); **F)** Motif of mouse ZFX identified by ChIP-seq; **G,H)** Motifs of human ZFY and mouse ZFY1 identified by Affinity-seq respectively.

Crystal structures of ZFPs bound with DNA show that most of the fingers interact with 3-4 base pairs (bp) and form tandem triplets^8–10^. Therefore, for a ZFP with N fingers, we would expect it to bind a sequence 3N bp long and have a motif of that length. Consistent with this, the mainstream motif prediction method predicts the full-length motif by concatenating the motifs for individual fingers derived from bacterial-one-hybrid (B1H) assays^11,12^. However, motifs identified for many ZFPs by ChIP-seq and HT-SELEX^13–15^ are significantly shorter than these predictions (Fig. 1B). To understand the source of this discrepancy, we used ZFY and CTCF as case examples to quantitatively characterize their specificity and learn the fundamental properties of tandem ZFP DNA recognition.

As the only zinc finger gene located on the human Y chromosome, ZFY has two close homologs in mice (mZFY1 and mZFY2), which are required for the meiotic sex chromosome inactivation (MSCI)^16^. ZFX, its counterpart at X chromosome, also shares close sequence homology. Via four complementary experimental approaches, we determined the DNA specificity for full-length ZFY and different truncated versions and assessed the contribution of different fingers to DNA binding affinity. We found that downstream fingers contribute to binding energy and confer specific recognition only in the presence of a perfect core site, which prompted us to propose a dependent recognition model different from the additive, position-independent recognition model. Using fluorescence anisotropy, we confirmed that the binding energy difference between various ZFY-DNA complexes is primarily driven by differences in dissociation rates, with a deviation from single-phase exponential curves that strongly suggested the existence of multi-mode recognition to its cognate sites.

The genome insulator CTCF was previously shown to confer multi-mode recognition to upstream, core, and downstream motifs by distinct sets of fingers, but the function of upstream and downstream fingers is unclear. In addition, while its sensitivity to methylated CpG (mCpG) within the core is critical for epigenetic regulation and imprinting control, the biophysical mechanisms underlying these roles remained not fully resolved^17–20^. First, by perturbing CTCF’s upstream and core sites simultaneously, we found that upstream specificity depends on the strength of the core, revealing another example of dependent recognition. Second, we quantified the energetic effect of various cytosine modifications (methylation, hydroxymethylation, formylation and carboxylation) within the core with different upstream sites and found that this effect depends on the strength of the upstream site. Structural comparisons between differently truncated CTCF-DNA complex revealed a small but significant structural tilt induced by the binding of upstream fingers, potentially explaining these results. Consistent with a critical *in vivo* role for dependent recognition, we further found that the mutant R567W - the first identified missense mutation in CTCF causing severe intellectual disability^21^ and possibly related to cancers^22^ - weakens the upstream recognition and therefore deregulates the epigenetic control within the core, providing a molecular mechanism for how mutations in accessory fingers indirectly affect genome-wide binding patterns and cause diseases.

These results from ZFY and CTCF suggested that dependent recognition between fingers could be a ubiquitous phenomenon for many more ZFPs. To test this hypothesis, we examined whether iterative motif analysis using a prefixed core help reveal longer motif profiles for ZNF343. Consistent with dependent recognition, auto-correlation and footprinting analysis of ChIP-exo data strongly suggested that only a small set of “good” cores facilitate upstream recognition. Overall, our work shows that current additive, position-independent recognition model is inaccurate to characterize the specificity and epigenetic property of tandem ZFPs and motivates a need for additional systematic, quantitative analysis to help us better understand how ZFPs work and how some mutants cause diseases.

## Results

### Short motifs are obtained for hZFY and mZFY1 using Affinity-seq

ZFY consists of an N-terminal activation domain followed by 13 tandem fingers (Fig. 1C); therefore, the mainstream motif prediction model suggests it should have a ~39bp long consensus binding site (Fig. 1D). Portions of this predicted motif are consistent with observed specificities for mouse ZFY orthologs bearing subsets of ZFY’s fingers with partially conserved amino acids (Fig. 1C): Taylor-Harris et al^23^ performed SELEX on mZFY1 and found it preferentially binds to sequences containing GGCCT, Grants et al^24^ showed that mZFY fingers 11-13 were sufficient to recognize a RGGCCT motif, and PBM^25^ and ChIP-seq^26^ experiments of ZFX produced the same 5 bp motif (Fig. 1E,F).

To identify preferred sequences for ZFY, we leveraged Affinity-seq, a method for *in vitro* selection of fragmented genomic DNA followed by MEME motif analysis^27^, to identify all bound sites from the entire genome sequence. Affinity-seq on human ZFY and mouse ZFY1 yielded 90,084 and 50,170 peaks at p values < 0.01, respectively, from which we found motifs very similar to those previously reported (Fig. 1G,H), with no secondary motifs reported by MEME.

### High-throughput SELEX (HT-SELEX) reveals a downstream consensus site and Spec-seq confirms this irregular downstream motif by fingers 7-11 of ZFY

To test whether ZFY has any extended motif beyond GGCCT, we adopted High-throughput SELEX (HT-SELEX)^13^, using randomized dsDNA libraries with a prefixed GGCCT in the flanking region (Fig. 2A). After two rounds of bound DNA selection by EMSA separation and amplification, we sequenced the enriched DNA pool and found the most enriched site to be GGCCT<u>AGGCGTTG</u>. We then fixed that extended consensus site, extended the randomized dsDNA region, and reran the SELEX assay, obtaining a most enriched site of GGCCTAGGCG<u>TTATTTT</u> (Fig. 2A).

**Figure 2.**
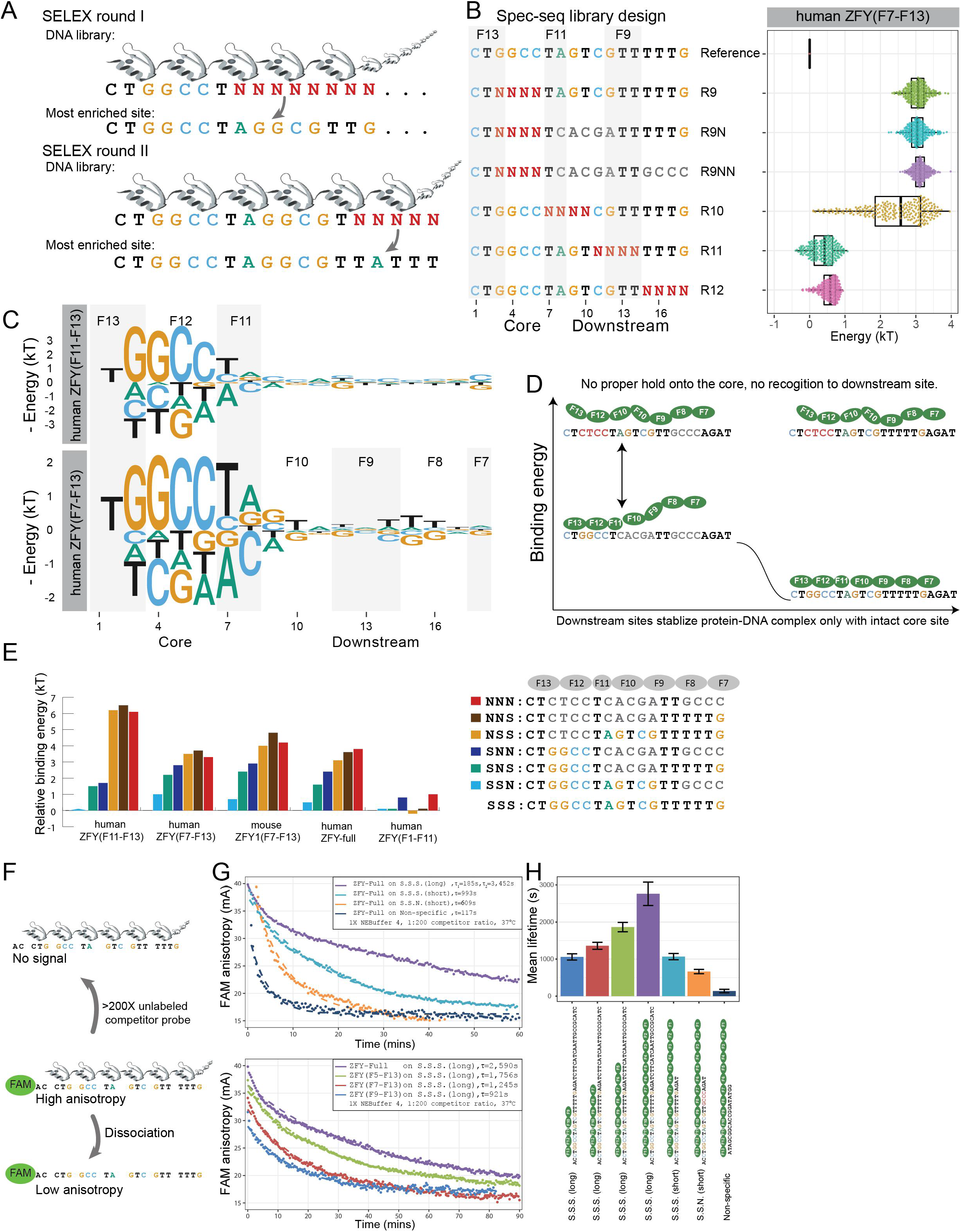
Quantitative analysis of ZFY reveals that its downstream recognition depends on the presence of perfect core. **A)** Workflow of two rounds of HT-SELEX with previously identified binding site as the anchor position; **B)** Spec-seq library design and the variants distribution of observed binding energy under human ZFY(F7-F13) construct; **C)** Position energy logo constructed by regression of all single variants of reference site; **D)** Dependent recognition model for Zinc Finger Y (ZFY); Without intact core, there is no observed specificity towards downstream region; **E)** Energy levels for representative binding sites under different constructs; S.S.N. is short name for Specific-Specific-Non-Specific, and so on; **F)** Workflow of florescence anisotropy assay to measure the intrinsic dissociation rates of various protein-DNA complexes; >200X unlabeled competitor probe was added to the pre-equilibrated binding reaction right before the kinetics monitoring processes; **G)** Upper panel shows dissociation curves for full-length ZFY over various DNA probes; Lower panel shows dissociation curves for full-length and various truncated ZFY constructs over S.S.S. (long) DNA probe; Single phase exponential curve was used to fit all observed data, except the ZFY-Full over S.S.S.(long), which shows two-phase exponential curve; **H)** Summary of observed dissociation rates of different constructs over various probes; Each sample was measured for at least three times; Sample means and standard deviations were derived from fitted single exponential curves.

Compared with other high-throughput techniques like HT-SELEX and Affinity-seq, Spec-seq^28^ is a medium-throughput method to quantitatively characterize the energy landscape of TF-DNA interactions with energetic resolution down to 0.2*k_B_T*. One practical limitation of Spec-seq is that one can only assay the relative binding energy up to a few thousands of variants, so having prior knowledge about the consensus site is preferable.

Using this SELEX-enriched site as a starting consensus sequence, we constructed tandem, non-overlapping dsDNA libraries for pilot Spec-seq runs and found that GGCCTAGTCGTTTTTG had slightly higher affinity than the SELEX-enriched site; therefore, we chose this sequence as the reference site for following runs. We designed four randomized dsDNA libraries (Rand 9,10,11,12) to tile across the entire reference site (Fig. 2B) and used Spec-seq to quantify relative binding energies for full-length mouse ZFY1, full-length human ZFY, and truncated versions of human ZFY containing different subset of ZFs (F11-F13, F9-F13, F7-F13, F5-F13, F1-F11). Consistent with previous work, we observed ^~^0.2*KT* measurement variation for individual sequences between replicate runs (Fig. S3), defining a practical resolution limit for significance.

From these Spec-seq results, we built position energy matrices (PEMs) and corresponding logos based on regression of the energy values of the chosen reference site and all its single variants (Fig. 2C). Compared to the hZFY(F11-F13), the motif for hZFY(F7-F13) revealed a downstream preference for a GT---TTT, indicating that those extra fingers contribute to this downstream motif. A comparison between logos for hZFY(F11-F13) and hZFY(F7-F13) reveals that finger 11 only recognizes a 2nt long TA motif in positions 7-8 rather than the predicted 3ntTCA in Fig. 1D. Visual comparison between B1H-predicted and ChIP-seq-discovered motifs suggests that this kind of irregular motif is quite common (e.g., as for ZNF140, ZNF324, and ZNF449 in Fig. S10).

### Recognition of downstream sites by ZFY requires an intact core

Next, we compared the binding energy of variants in the Rand9 library (R9) to two additional libraries (R9N, R9NN) containing the same randomized sequences within the core but some mismatches downstream. All variants within the R9, R9N, R9NN libraries were approximately 3kT higher, with no difference in median binding energy between libraries, suggesting that any single mismatch within the core is sufficient to abolish specific recognition (Fig. 2B). For sites in the non-specific plateau with defective cores, there is no practical value to predict their energy using PEMs derived from single variant data (Fig. S4), and presence of downstream motifs does not contribute extra binding energy (Fig. 2D).

To further probe this unexpected non-additivity, we quantified binding of various full-length and truncated constructs (hZFY(F1-F11), hZFY(F11-F13), hZFY(F7-F13), mZFY1(F7-F13) and full-length hZFY) to a few representative sequences (Fig. 2E). For example, the site CTGGCTAGTCGTTGCCC contains a specific core, a specific downstream site at position 7-12, and a non-specific downstream site at 13-18, which we designate as Specific-Specific-Non-specific, or SSN for short. While hZFY(F11-F13), hZFY(F7-F13), mZFY1(F7-F13) and hZFY-full clearly show similar relative patterns of recognition to sites containing an intact core (*i.e*. the SSS reference sequence is bound more strongly than SSN, SNS, and SNN sites due to preferential recognition by downstream fingers), there is no consistent difference between measured energies for NXX sites (*i.e.* NSS, NNS, and NNN). In addition, measured binding for the hZFY(F1-F11) mutant, which lacks fingers required for core site recognition, reveals no significant energetic differences between sequences (Fig. 2E). Together, these measurements demonstrate that specific recognition of downstream sequences depends on core binding in the first place.

### ZFY-DNA dissociation rates correlate with affinities and reveal alternative modes of recognition

In addition to energy measurements, we directly measured the intrinsic dissociation rates *k_off_* for various ZFY constructs interacting with different DNA sequences by fluorescence anisotropy. After binding reactions reached equilibrium, we added a 200-fold excess of unlabeled competitor DNA and monitored anisotropy values of the FAM-DNA probes over time to visualize dissociation processes (Fig. 2F, 2G). Changes in mean lifetime for full-length hZFY interacting with different DNA sequences were consistent with Spec-seq measurements: SSS and SSN had a 0.7*kT* energy or 2-fold affinity difference as measured by Spec-seq, and their mean lifetimes also differed by 2-fold (1,176s and 656s, respectively), suggesting that measured energy differences for ZFY-DNA binding are primarily driven by differences in their dissociation rates. However, while measured dissociation curves for truncated ZFYs interacting with SSS(long) probe and full-length ZFY interacting with SSN (short) or nonspecific probes were well-fit by a single exponential curve, full-length ZFY dissociation from SSS (long) probe showed significant deviation from single exponential behavior (Fig. 2G lower panel). We resorted to two-phase exponential curve fitting, which yielded better results (Fig. 2G, upper panel). These observations strongly suggest that a simple two-state DNA binding model is inadequate to address the complex, multi-mode recognition of ZFP to extended, non-additive motifs.

### Upstream specificity profile of CTCF depends on the strength of the core

Next, we explored whether other long ZFPs also show significant non-additivity in binding site recognition. CTCF, the genome insulator in the human genome, is composed of 11 tandem zinc fingers that have identical sequences between humans and mice (Fig. 3A). Previous ChIP-chip and ChIP-seq work^14,29,30^ identified a 14nt core motif CCNNNAGGGGGCGC recognized by fingers 7 to 3. Later Nakahashi, H. *et al*.^31^ reported extra upstream and downstream motifs with a variable 5-6 nt distance to the core (Fig. 3A, downstream motif not included). According to their analysis, within 48,137 detected ChIP-seq binding peaks, all CTCF binding sites contained a motif feature matching the core, but only a subset of these (^~^6,000 sites) contained flanking sequences matching the upstream motif adjacent to the cores. If CTCF binding site recognition were truly independent, we would expect observed ChIP-seq peaks to also contain sites matching the upstream motif alone in the absence of the core; however, this was not observed.

**Figure 3.**
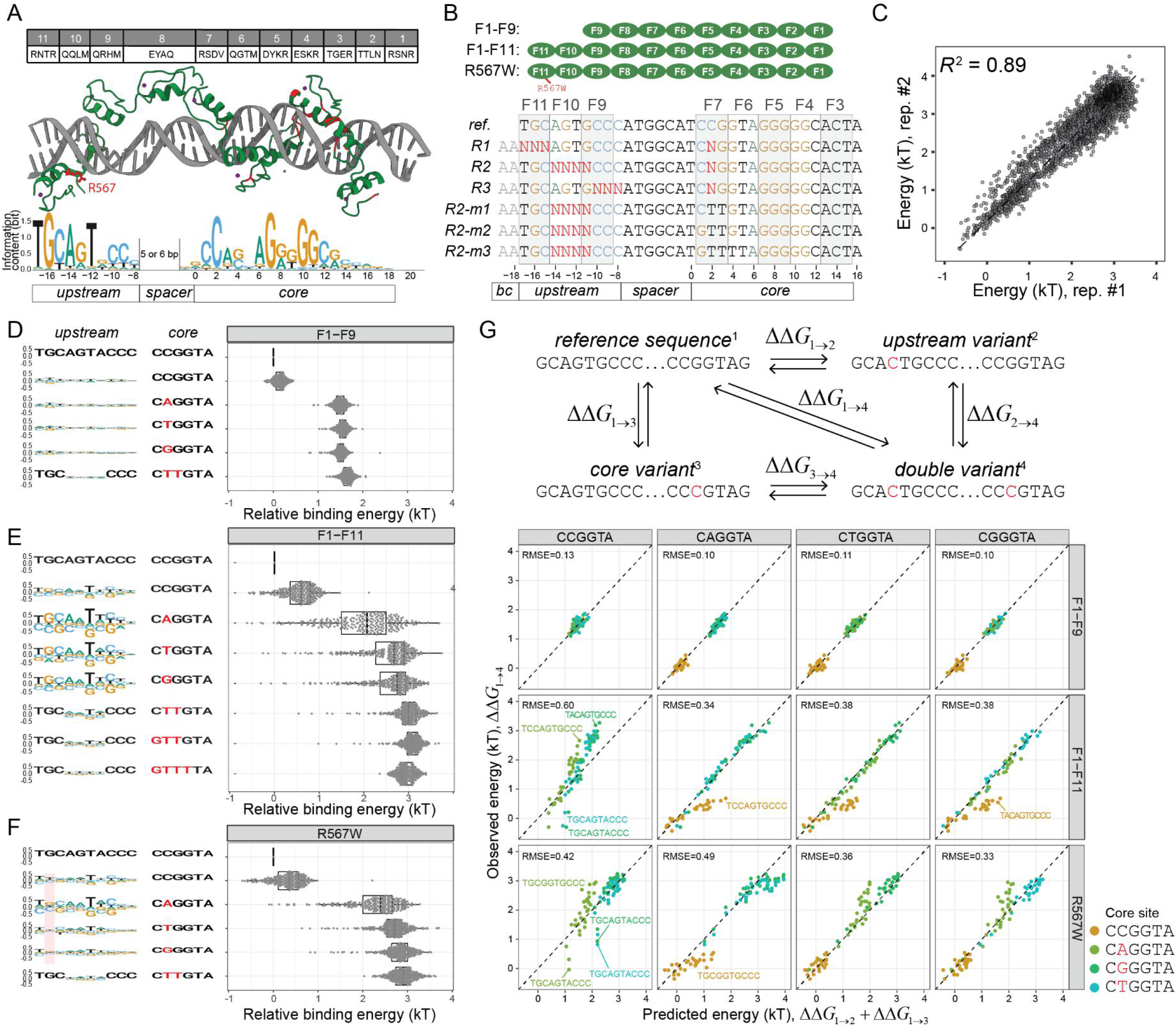
The upstream specificity of CTCF depends on the strength of core sites and violates additivity assumption. **A)** Contact residues composition for human/mouse CTCF and current structural model about CTCF’s recognition to its binding site; Currently identified missense disease mutants are mapped to the structure and labeled red; The upstream and core motifs are based on previous ChIP-exo results; **B)** CTCF constructs used in current study (N-terminal HALO-tag not shown); Spec-seq libraries design for unmethylated sites; R1, R2, and R3 randomize the core and upstream sites simultaneously, whereas R2-m1, m2, and m3 carry defective cores along with randomized upstream sequences; **C)** Spec-seq data reproducibility between replicates; All observed energy values are normalized against the reference site in each sample; **D, E, F)** Energy distribution of variants in R2 libraries with different cores are shown on the right panel, whereas the upstream motifs derived based on the corresponding cores are shown on the left panel; Part of the R567W motif that differs from wildtype are highlighted in red; **G)** Additivity test for CTCF using four reference sites with different cores (CCGGTA, CAGGTA, CGGGTA, and CTGGTA); The observed value of double variant is plotted against the predicted value of double variant based on single variant data; R.M.S.E. were shown for each case.

To test if CTCF recognition is additive and how upstream specificity changes with different cores, we designed Spec-seq libraries (Fig. 3B) in which we simultaneously altered the upstream and core sequences (R1, R2, R3). In addition, we designed three libraries (R2-m1, R2-m2, and R2-m3) to incrementally introduce more mismatches into the core and thereby test if the upstream site can still be properly recognized by fingers 9-11 in the presence of increasingly defective cores. Overall, we profiled binding of three CTCF constructs (F1-F9, F1-F11, and the known disease mutant R567W) to each library and observed good reproducibility between replicates (Fig. 3C, S3).

For each construct, we then sorted results by the strength of the core and generated a logo to depict the observed upstream specificity. Consistent with previous results, without finger 10 and 11, the truncated F1-F9 construct shows no upstream specificity at all (Fig. 3D), while for the F1-F11 construct, the upstream site TGCAATCCC was the optimal site associated with most cores (Fig. 3E). The measured energy distribution of upstream variants varied significantly from core to core, with the strongest core CCGGT showing modest upstream specificity and cores of intermediate strength (C<u>T</u>GGT, C<u>G</u>GGT, C<u>A</u>GGT) exhibiting the biggest dynamic range (up to 4kT) and the strongest upstream specificity. For highly defective cores like <u>GTTTT</u>A, we didn’t observe any upstream motif. These results are consistent with a model in which upstream sites need only contribute a small amount of energy to form specific, stable protein-DNA complex in the presence of the strongest core, but even the strongest upstream site alone is insufficient to localize CTCF to a very weak core. These observations again cannot be explained by an additive, position-independent recognition model, under which the upstream motif profile should have no correlation with core strength at all.

We then attempted to directly quantify non-additivity via double mutant cycle analysis, in which we chose four different reference sites (CCGGT, CAGGT, CTGGT, CGGGT), perturbed the upstream and core each with single mismatches alone and in combination, and compared the observed energy of double variants to what would be predicted assuming additivity of single variant effects (Fig. 3G, upper panel). While double variants at some sites contributed additively, others showed significant non-additivity, particularly when using the strong core CCGGT as reference (Fig. 3G, lower panel). This observed non-additivity has practical consequences for motif finding: for example, using the weak upstream motif observed in the presence of the strong core CCGGT to predict binding energies for other sites with weaker cores (CAGGT, CTGGT, CGGGT) will systematically underestimate the true binding energy.

Besides the regular R2 library in our design, we included an ‘R2L’ library to test whether CTCF can recognize the upstream motif in extended 6nt spacing format, as previously reported. Indeed, CTCF was still able to recognize the upstream site with the extended spacing format, though the observed motif appeared slightly weaker than the regular spacing case (Fig. S5). Together, these results establish that CTCF binds DNA non-additively with significant implications for motif discovery: we cannot reliably predict the binding energy for all sites with one single PWM or PEM for long ZFPs.

### Upstream sites negatively regulate effects of cytosine modifications within the core

The methylation effect, or C-to-mC substitution effect, can be defined as the energy difference between methylated and unmethylated sites sharing the same sequence such that a positive effect means that methylation blocks protein-DNA interactions, known to be important for epigenetic control by CTCF^17^. Previously, our scanning of CTCF’s core site revealed that only mCpG at position 2 to 3 confers a significant energy change^32^, later shown to be exclusively derived from an upper strand mC at position 2 recognized by finger 7^18^. However, to date there has been no systematic study of the effect of other epigenetic modifications such as hydroxymethylation, formylation, and carboxylation (Fig. 4A).

**Figure 4.**
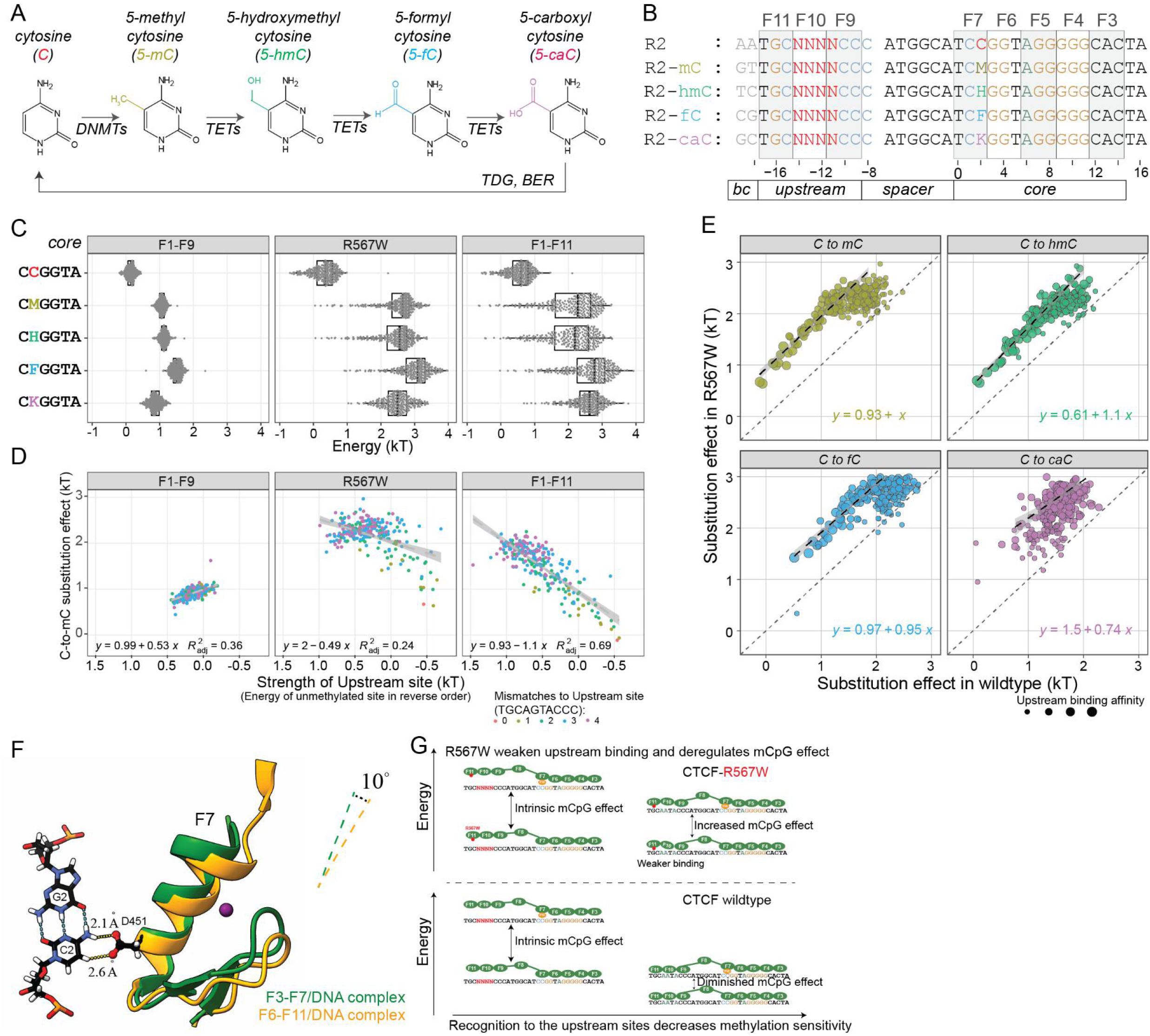
The methylation effect of CTCF recognition is negatively regulated by the strength of upstream sites. **A)** Known cytosine methylation and modification pathways in human; **B)** Methyl-Spec-seq libraries design testing various C-to-xC effects at position 2 with altered upstream sites; Barcodes at −19 and −18 indicate the type of modifications; **C)** Variants distribution of binding energy with different modifications and constructs; M, H, F, K are short for methylated, hemimethylated, formyl, and carboxyl cytosines respectively; **D)** Relationship between observed C-to-mC substitution effects and the Strength of Upstream site, which is defined by the energy of corresponding unmethylated site; **E)** Comparison of C-to-xC substitution effects between wildtype and R567W mutant constructs; The dashed lines indicate linear regression of high affinity sites; **F)** Structural comparison of existing CTCF-DNA complexes; Two structures were aligned over cytosine base at position 2; Only CG at position and Finger 7 were shown; **G)** Proposed Upstream Regulation model of CTCF.

To test whether these other modifications affect the binding affinity of CTCF and whether variations in the upstream site affect the magnitude of observed methylation effects, we applied our previously developed method Methyl-Spec-seq^32^. These measurements used a revised Spec-seq libraries design that included four more randomized libraries (R2-mC, R2-hmC, R2-fC, R2-caC) carrying chemically modified cytosines at the upper strand of position 2 as designated and modification-specific barcodes at positions −19 to −18 (Fig. 4B).

We initially expected that any modification sufficient to disrupt finger 7 recognition would also abolish upstream site recognition, thereby enhancing the observed epigenetic effect in the presence of a strong upstream site. However, we observed several surprising results. While all tested cytosine modifications at position 2 blocked DNA binding of CTCF to different degrees (Fig. 4C), TGCAATACCC always remained the optimal upstream site, suggesting that a single chemical modification is far from disruptive enough to abolish the upstream recognition. Moreover, plotting the C-to-mC effect *vs.* the strength of the upstream site (calculated from the unmethylated case) revealed a negative correlation for the F1-F11 full-length construct but no correlation for the truncated F1-F9 construct (Fig. 4D). For the optimal upstream site TGCAATACCC, the C-to-mC effect is even diminished (Fig. S6). This result was consistent across replicates and experiments and across multiple modifications (hmC, fC, and caC) (Fig. S6).

Currently the C-to-mC effect of CTCF is thought to be caused by the steric clash of aspartate residue 451 (D451) with the methylated cytosine at position 2 (C2)^18^. While there is not yet a full-length CTCF-DNA crystal structure, we investigated the structural relationship between CTCF and the DNA interface by aligning partially overlapping CTCF-DNA structures as surrogates (PDB #5KKQ and #5YEL) based on the C2 nucleotide and comparing their orientation difference (Fig. 4F). These aligned structures show that the recognition helix of finger 7 is tilted 10 degrees away from C2 in the presence of upstream recognition, prompting us to propose a “Grip-and-Control” model (Fig. 4G) in which fingers 9-11 properly ‘grip’ the upstream sites and form a specific complex, leading to a conformational shift that yields a CTCF-DNA complex that is more tolerant to internal single mismatches and chemical modifications, thereby decreasing the observed C-to-mC effect. Note that the carboxyl group is bulkier, rendering its effect under upstream control weaker than other modifications (Fig. S6).

From a functional perspective, this means that the epigenetic effect of methylating each individual CTCF binding site in the human genome can be quantitatively fine-tuned by the flanking upstream sequence to the desired level (0 to 2kT), which could be important to fulfill the regulatory function of CTCF.

### Mutant R567W in finger 11 weakens upstream recognition and deregulates the methylation effect to the core

Next, we constructed a full-length CTCF structural model by aligning published structures^18,33^, and mapping all currently identified disease mutants^34^ onto this model (Fig. 3A). Remarkably, all missense mutations are located within base-touching fingers (Fig. 3A). R567W is particularly interesting, as it is the first identified missense mutant, originally found to cause intellectual disability, microcephaly, and growth retardation^21^, and was later found in endometrial endometrioid adenocarcinoma, endometrial mixed adenocarcinoma, and lung adenocarcinoma^22^. While R567W’s location in finger 11’s recognition helix opposing the base-contacting side suggested this mutant should only marginally affect upstream recognition while having no influence over the core, its surprising association with dementia more severe than that associated with nonsense and other missense mutants^21,35^ led us to characterize it more deeply.

Methyl-Spec-seq results for R567W showed two differences from wildtype CTCF. First, R567W altered upstream specificity mainly in the R1 region (position −17 to −15, Fig. 3F), altering the preferred motif from TGC to TtC. The existing structure shows arginine 567 loosely associated with the phosphate backbone, suggesting that mutating this residue could weaken the local recognition even without directly touching the base. Second, if we view the upstream fingers as a regulatory module, weakening this module could deregulate and increase the C-to-mC effect to the core. For all binding sites tested in our libraries (including high affinity upstream sequences and all different methyl modifications), the modification effect at C2 for R567W was almost always bigger than for the WT case (Fig. 4E). *In vivo*, we expect that this R567W-mediated deregulation could confer higher-than-normal methylation sensitivity and lower-than-normal occupancy for CTCF sites with mCpG or mismatches within the core. This may partly explain the observed clinical severity of R567W mutations: while low CTCF occupancy due to haploinsufficiency could be compensated by higher CTCF expression levels, epigenetic defects might be more difficult to rescue.

### Iterative analysis of ZNF343 ChIP-exo data reveals dependent recognition relationship between upstream and core

These results for ZFY and CTCF suggest that dependent recognition, in which ZFPs cannot properly recognize flanking upstream/downstream sites in the vicinity of defective cores, could be a general property for other ZFPs. As a specific example, human ZNF343 is a 12-finger long KRAB-ZFP, yet RCADE analysis^15^ of published ChIP-exo data^36^ only reveals a 6-nt long motif (GAAGCG) (Fig. 5C). The B1H prediction model (Fig. 5B) suggests that this hexamer site is most likely recognized by fingers 2 to 1. With this prior knowledge, we identified 3,237 GAAGCG sites within the reported 4,532 ChIP-exo peaks and then used these sites as a fiducial reference to align, map, and count all ChIP-exo reads based on their relative distance to the nearest GAAGCG site (Fig. 5F). The aggregate ChIP-exo traces revealed a significant reverse peak signal at position 7, suggesting that this hexamer is indeed recognized by fingers 2-1. Ideally, the ChIP-exo read counts near each site should be proportional to the binding occupancy; if the *in vivo* ZNF343 protein concentration is low enough, reads should also be proportional to the binding affinity. Based on these assumptions, we calculated the negative logarithmic ratio of ChIP-exo reads near each site as an estimate of the relative binding energy, and then performed data regression and motif analysis (Fig. 5D). This iterative analysis revealed an upstream motif in positions −8 to 0 that is very similar to the published result of HT-SELEX^37^ (Fig 5E).

**Figure 5.**
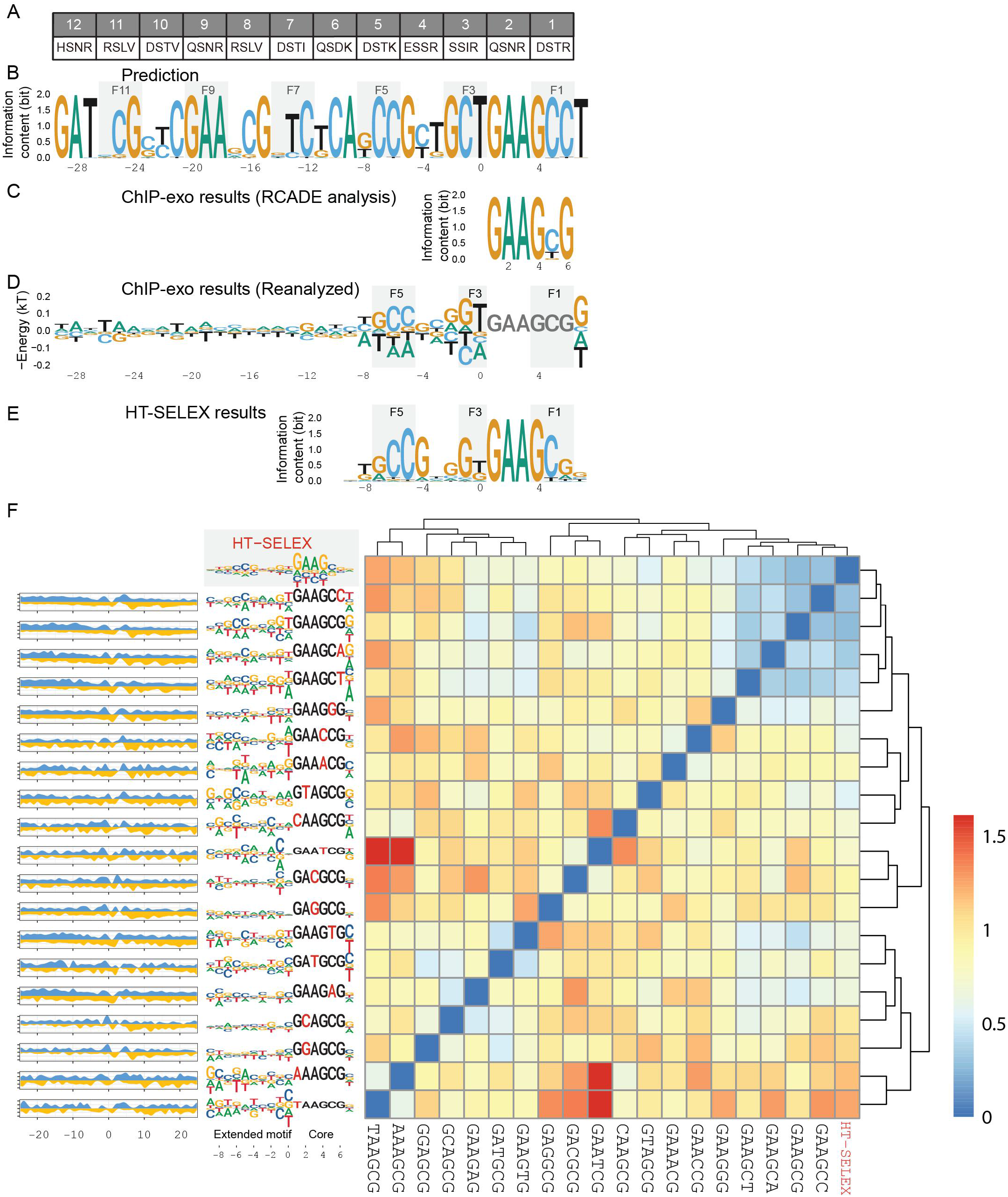
Iterative motif analysis of ZNF343 reveals the dependent relationship between core and upstream motifs. **A)** Contact residues for human ZNF343; **B)** Motif prediction by B1H method; **C)** Motif from RCADE analysis of ChIP-exo data; **D)** Extended motif by reanalysis of ChIP-exo data with prefixed core GAAGCG; **E)** HT-SELEX results of ZNF343; **F)** Extended motifs by reanalysis of ChIP-exo data with all single variants of GAAGCG as the prefixed core; Heatmap is generated by auto-correlation analysis of all extended motifs with different cores and HT-SELEX result; The ChIP-exo reads footprints near associated prefixed cores are shown on the left.

Encouraged by these results, we then applied a similar iterative analysis to all other cores with no more than one mismatch to GAAGCG and performed hierarchical clustering and auto-correlation analysis based on similarity of the resulting upstream motifs to each other (Fig. 5F). This analysis reveals that only those cores differing with GAAGCG at peripheral position 6 can serve as “good” cores to facilitate upstream recognition, and that internally mismatched cores at position 1-5 severely compromise upstream specificity. Indeed, the HT-SELEX result shows that position 6 has a relatively weak motif and a limited contribution to the core. Also, only the ChIP-exo footprints of these good cores show the expected asymmetric patterns we observed in the intact case. To facilitate broad adoption by other labs to study additional multi-domain ZFPs and reveal dependent recognition, we have deposited this data analysis protocol in a public repository.

## Discussion

To tackle the “long fingers but short motifs” paradox, we must first ask: is it true that long ZFPs really have short motifs? We argue that there are two technical issues that hinder long motif discovery and a biophysical reasoning for why PWMs should not be used blindly for specificity modelling. First, most existing techniques like PBM, ChIP-seq, and Affinity-seq have limited power to resolve weak binding sites. For ChIP-seq measurements of occupancy on human genomic DNA, each 15-mer will show up only once on average, making the discovery of longer motifs increasingly difficult. Second, the irregular nature of extended motifs like those seen for ZFY, where each finger does not appear to interact with the ‘expected’ three bases, can also inhibit motif detection when searches are guided by motif prediction methods. A visual comparison between the predicted motifs and ChIP-seq results for 131 human ZFPs with reported motifs suggests that such irregular motifs may be quite common: for example, ZNF140, ZNF324, and ZNF449 all likely contain some irregular motifs with fewer than three base-pairs for some internal fingers (Fig. S10). Lastly, but most importantly, when we talk about motifs, we typically refer to some additive, position-independent recognition model like a position weight matrix (PWM), which assumes each base contributes to the overall binding energy additively and independently from bases in other positions. While such models have shown broad utility for a wide variety of TFs across many structural classes, there was never a thorough, quantitative analysis to justify the validity of these assumptions for tandem ZFPs. Here, we showed that proper recognition to a “good enough” core must precede recognition of flanking sites and that the specificity in the core is modulated by the flanking sequence. It is fair to say that a position-independent weight matrix is inaccurate and inappropriate to characterize the specificity profile of tandem ZFPs.

In future work, improved motif discovery algorithms should consider these intrinsic properties of long ZFPs, including irregular motif structure, variable spacing and dependent recognition between sub-motifs. Because of the dependence of flanking motifs on the core, it is computationally more effective to identify the core motif first and use it as anchor site to analyze flanking regions to extract more motif information for other ZFPs, as we did for ZNF343. Moreover, visualization techniques that explicitly depict nucleotide interdependence^38^ and Contribution Weight Matrices (CWM)^39^ can help reveal these non-additivities.

If long ZFPs do have long motifs, what are the functions of those extra fingers? For CTCF, while most of its *in vivo* binding sites seem not to have flanking upstream or downstream sites, comparative genomic analysis shows that all eleven fingers are highly conserved across mammals. Here, we show that those extra fingers can modulate the specificity and methylation sensitivity to the core by tuning the strength of upstream sites, essentially serving as an “epigenetic modulator”. This may not be unique: many KRAB-family ZFPs have exceedingly long arrays of fingers generally thought to be engaged in silencing of transposable elements (TEs). Those extra fingers may be required to facilitate a variety of additional functions including preferential recognition of TEs over non-TEs, worthy of further investigation.

## Experimental and Data analysis Procedures

### Construction and expression of recombinant proteins

The coding sequences for human ZFY (Uniprot P08048:408-768) and mouse ZFY1 (Unirprot P10925:390-782) were codon optimized for E. coli expression and synthesized as IDT gBlocks and mouse CTCF (Uniprot Q61164-1:241-583) was cloned from mouse cDNA libraries. After In-Fusion cloning into an NEB DHFR control vector with an N-terminal hisSUMO tag, proteins were expressed and purified largely as in our previous work except that we used an extra heparin column purification to increase purity for anisotropy experiments. For Methyl-Spec-seq experiments of CTCF, we used the NEB PURExpress system to produce N-terminal HALO tagged CTCF constructs and noticed better success rates than previous SUMO-tagged constructs. All constructs, including truncated versions, are listed in Supplemental Table S1.

### Affinity-seq procedures

Affinity-seq was essentially done as in [36] with minor adjustments. A ZF array of the protein of interest was amplified then cloned into a universal Affinity-seq vector by recombineering. The resulting construct expressed a fused protein containing 6HisHALO–the 412-511 aa fragment of PRDM9–ZF array of interest. The fused protein was expressed in Rosetta 2 cells at 15°C for 24 h and partially purified by ion exchange chromatography on SP-sepharose. The purified protein was mixed with genomic DNA sheared to ^~^200 bp on a Covaris ultrasonicator, and allowed to bind overnight. The protein-DNA complexes were then isolated on HisPur Ni-NTA Resin (Thermo Scientific) preincubated with a partially purified prep of the empty tag to reduce the background. DNA was then eluted and used to prepare genomic libraries using a TruSeq ChIP Library Prep Kit (Illumina). The libraries were sequenced on a HiSeq2500 or NextSeq platform ensuring ^~^50 million reads per library. Data were analyzed using a custom pipeline as described previously [36] and we performed motif analysis using the MEME software package.

### HT-SELEX procedures

For the first round of HT-SELEX, ^~^200ng dsDNA libraries containing randomized sequence CAGGCCTNNNNNNNN were used for EMSA shift with hisSUMO-hZFY titrated from low to high concentration. Each time, only the lane containing the lowest amount of protein was chosen and the bound portion of DNA (no more than 20% of total DNA) was cut and then amplified for the next round of SELEX selection enrichment. Since in the first round of HT-SELEX, the most enriched site turned out to be CAGGCCTAGGCGTTG, the DNA library was redesigned as CAGGCCTAGGCGTNNNNNNNN for further HT-SELEX by EMSA separation. Again, each time we ensured that no more than 20% of the total DNA was in the bound state for selection and enrichment analysis.

### Spec-seq, Methyl-Spec-seq procedures and motif analysis

The experimental procedures were essentially the same as our previous work, with all binding reactions set up at 1X NEBuffer 4, room temperature. For ZFY, EMSAs were performed using 9% Tris-glycine gels in the cold room run at 200V for 30mins. We noticed that for some ZFY proteins, particularly ZFY(F11-F13), when the protein concentration was too high, the shifted DNA fragments appeared easily to form protein oligomers or aggregate near the EMSA well. Consequently, we generally used low concentrations of protein (<100nM) and selected only monomeric ZFY-DNA complexes for Spec-seq analysis. For CTCF, 10% Tris-glycine gels were used to separate the bound and unbound fractions of DNA (again in the cold room and run at 200V for 50mins). Position energy matrices or energy motifs were derived by data regression of the binding energy of either reference sites plus single variants or all measured sites using the TFCookbook package.

### Dissociation kinetics assay by fluorescence anisotropy

All binding assays were performed in 1X NEBuffer 4 at 37°C with 30nM FAM-DNA probe, and in this condition the basal value for FAM-DNA probe without protein was ^~^15mA. With a saturating concentration of ZFY protein added, the anisotropy values can go above 100mA. In our case, we titrated a low volume of protein (<4% v/v), yielding initial values at equilibrium only above 40mA, suggesting that only a small fraction of the DNA was bound (<20%) and the DNA probe was more likely bound by the protein in an assumed specific conformation. After we injected highly concentrated unlabeled competitor DNA (500pM/uL X 2uL) into 100uL binding reactions, yielding a molar ratio between FAM probe and competitor DNA < 1:200, we measured anisotropy values at 20s or 40s time intervals for up to 90mins.

To measure the intrinsic dissociation rates of ZFY-DNA complexes, we did titration experiments first with different molar ratios of unlabeled competitor DNA into the binding reactions, as in Fig. S7. For competition ratios below 1:100, the observed dissociation rate reached some plateau and did not increase further. Therefore, we assumed it was appropriate to use the 1:200 competitor ratio curve to estimate the intrinsic dissociation rate or mean lifetime of the protein-DNA complex.

After setting up the binding reaction for at least 20mins, we assumed the system had reached equilibrium state. Slightly to our surprise, however, we observed the anisotropy value slowly decrease over time even in the absence of any added competitor DNA, likely representing steady inactivation or degradation of ZFY protein at 37C. To exclude the possibility that the measured dissociation rates are differentially biased by different protein inactivation rates, we measured this inactivation process alone for different proteins, and they all showed very similar inactivation rates, which are significantly slower than our observed dissociation rates (Fig. S8).

To quantify the dissociation rate k_off_ or mean lifetime τ, we fit our data using single exponential decay model with following equation:

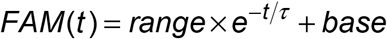

The base value is usually in the range 15 to 17, and the range parameter depends on the first measured anisotropy value in each experiment, which should not affect the mean lifetime. For full-length ZFY construct interacting with the S.S.S.(long) probe, we noticed a significant discrepancy between observed data and fitted curves, so a two-phase exponential decay model was also used:

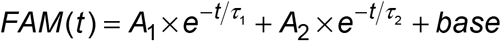

Each experiment was repeated at least three times to calculate the mean values and standard deviations (as in Table S2). All experiments were performed using a TECAN Safire2 instrument set to 490nm excitation/525nm emission wavelength.

### Motif analysis of human ZNF343 using published ChIP-exo data

The detailed analysis workflow and processed data of ZNF343 are deposited in GitHub (https://github.com/zeropin/ZNF343-analysis-workflow) for free re-use.

### Alignment and comparison of published CTCF structure models

ChimeraX was used to superimpose two structure models (PDB #5KKQ, #5YEL) based on alignment of the cytosine at position 2.

## Supplemental Information and Data availability

Supplemental information Includes all nucleotide sequences, experimental conditions and procedures, anisotropy measurement descriptions and record, and irregular motif predictions. All raw and processed sequencing data were deposited to NCBI GEO database (Affinity-seq #GSE111772, ZFY Spec-seq #GSE109098, CTCF Spec-seq #GSE110155). The software package used for motif regression and analysis can be accessed through (https://github.com/zeropin/zeropin/TFCookbook).

## Author Contributions

Z.Z. conceived this project, designed and performed the HT-SELEX, Spec-seq and anisotropy dissociation experiments in G.D.S. lab, designed and performed Methyl-Spec-seq in P.M.F. lab, analyzed ZNF343 data, wrote the first draft. P.M.P., T.B., and M.W. performed and analyzed the Affinity-seq experiments. G.D.S. supervised this work. P.M.F. supervised this work and co-wrote the manuscript.

## Acknowledgement and Funding

We thank Dr. Rafael Casellas for providing the CTCF PWM data for motif comparison purposes. We also thank anonymous reviewers of the former manuscript for helpful suggestions to improve the presentation. This work was supported by NIH grants HG000249 (GDS), GM078452 (PMP) and 1DP2GM123641-01 (PMF). P.M.F. is a Chan Zuckerberg Biohub Investigator.

